# Tissue-resident alveolar macrophages reduce O_3_-induced inflammation via MerTK mediated efferocytosis

**DOI:** 10.1101/2023.11.06.565865

**Authors:** M.A. Guttenberg, A.T. Vose, A. Birukova, K. Lewars, R.I. Cumming, M.C. Albright, J.I. Mark, C.J. Salazar, S. Swaminathan, Z. Yu, Yu. V. Sokolenko, E. Bunyan, M.J. Yaeger, M.B. Fessler, L.G. Que, K.M. Gowdy, A.V. Misharin, R.M. Tighe

## Abstract

Lung inflammation, caused by acute exposure to ozone (O_3_) – one of the six criteria air pollutants – is a significant source of morbidity in susceptible individuals. Alveolar macrophages (AMØs) are the most abundant immune cells in the normal lung and their number increases following O_3_ exposure. However, the role of AMØs in promoting or limiting O_3_-induced lung inflammation has not been clearly defined. Here, we used a mouse model of acute O_3_ exposure, lineage tracing, genetic knockouts, and data from O_3_-exposed human volunteers to define the role and ontogeny of AMØs during acute O_3_ exposure. Lineage tracing experiments showed that 12, 24, and 72 h after exposure to O_3_ (2 ppm) for 3h all AMØs were tissue-resident origin. Similarly, in humans exposed to FA and O_3_ (200 ppb) for 135 minutes, we did not observe ∼21h post-exposure an increase in monocyte-derived AMØs by flow cytometry. Highlighting a role for tissue-resident AMØs, we demonstrate that depletion of tissue-resident AMØs with clodronate-loaded liposomes led to persistence of neutrophils in the alveolar space after O_3_ exposure, suggesting that impaired neutrophil clearance (i.e., efferocytosis) leads to prolonged lung inflammation. Moreover, depletion of tissue-resident AMØ demonstrated reduced clearance of intratracheally instilled apoptotic Jurkat cells, consistent with reduced efferocytosis. Genetic ablation of MerTK – a key receptor involved in efferocytosis – also resulted in impaired clearance of apoptotic neutrophils followed O_3_ exposure. Overall, these findings underscore the pivotal role of tissue-resident AMØs in resolving O_3_-induced inflammation via MerTK-mediated efferocytosis.

## Introduction

Air pollution is responsible for a significant number of premature deaths globally with an average mortality of 6.5 million individuals per year.^1^ Despite an already significant impact on rates of global morbidity and mortality, air pollution associated mortality will continue to rise with climate change. Furthermore, current regulatory efforts will not eliminate air pollution-associated morbidity and mortality, particularly in sensitive individuals.^1,2^ Understanding the mechanisms that drive air pollution-mediated health effects is therefore critical to better target mechanisms that mitigate its adverse effects.

When ground-level ozone (O_3_) exposures increase, as with other criterion air pollutants, the risk of respiratory-related morbidity and mortality increases.^3–5^ A principal lung effect of O_3_ exposure is the generation of lung inflammation and injury. Generally, O_3_-induced lung inflammation and injury is considered mild and resolving; however, in sensitive individuals or those with underlying lung disease this can lead to exacerbated or prolonged inflammation and injury responses.^6^ Lung inflammation is a highly regulated process that includes both initiation and resolution phases that are regulated, in part, by signaling of alveolar macrophages (AMØ).^3,7^ Prior research underscores the critical roles of AMØs in O_3_-induced lung injury. This includes well-characterized pro-inflammatory effects including cytokine generation^8^, and airspace neutrophil influx.^9^ Conversely, AMØs can function in the resolution of lung inflammation via functions such as phagocytosis and efferocytosis that clear pathogens, cellular debris, and apoptotic immune cells.^10^ This diversity of AMØs functions has made them a challenge to study and has limited efforts to target them therapeutically. Thus, a better understanding of the context-dependent functions of AMØs following O_3_ exposure are required to target them with precision.

AMØ functions can depend on their specific origin (i.e., ontogeny). In naïve mice, all AMØs are of tissue-resident origin. These tissue-resident AMØs populate the lung during early postnatal period and, in the absence of lung injury, maintain themselves via proliferation *in situ* for the life of the animal.^11,12^ Alternatively, following lung injury, monocytes are recruited to the lung and differentiate into monocyte-derived AMØ.^13,14^ Though tissue-resident and monocyte-derived AMØs share some common macrophage functions, they also exhibit distinct metabolism, activation markers, and cytokine profiles suggesting different functional roles in lung injury and repair.^15^ While monocyte-derived AMØs have clearly documented roles in promoting lung injury, inflammation, and fibrosis,^15–17^ the role of tissue-resident AMØs is less understood. Furthermore, the composition and function of AMØs following the mild and resolving acute lung injury with acute O_3_ exposure has not been clearly defined.

To address this question, we defined the composition of AMØs following filtered air (FA) and O_3_ exposure using a lineage-tracing mouse system which labels monocytes and monocyte-derived AMØs but not tissue-resident AMØs.^17,18^ Following O_3_ exposure in these mice, we did not identify an increase monocyte-derived AMØs. Additionally, in healthy human volunteers undergoing laboratory FA and O_3_ exposure, we also did not observe an increase in monocyte-derived AMØs. To define tissue-resident AMØ functions in O_3_-induced lung injury, we administered intratracheal clodronate-loaded liposomes to deplete tissue-resident AMØs^16^ and then exposed them to FA or O_3_. Depletion of tissue-resident AMØs resulted in persistence of O_3_-induced airspace neutrophilia, suggesting a defect in clearance of apoptotic neutrophils (i.e., efferocytosis). We confirmed the direct effect of tissue-resident AMØs on efferocytosis by demonstrating reduced clearance of apoptotic Jurkat cells following clodronate-mediated depletion of tissue-resident AMØ. Finally, in MerTK^-/-^ mice, which have defective efferocytosis ^19,20^, we defined the impact of reduced efferocytosis on O_3_-induced lung inflammation.

Consistent with tissue-resident AMØ-depleted mice, O_3_-exposed MerTK^-/-^ mice exhibited persistent airspace neutrophil inflammation. These data support that tissue-resident AMØs promote resolution of O_3_-induced lung inflammation via MerTK-mediated efferocytosis of neutrophils.

## Materials and Methods

### Experimental Animals

C57Bl/6J mice were purchased from Jackson Laboratories. The mice were a mix of males and females, group specific composition listed in figure legends. MerTK^-/-^ and Cx3cr1^ERcre^ zsGreen mice were bred in-house. MerTK^-/-^ mice were a kind gift from Dr. Edward Thorp, Northwestern University; originally from The Jackson Laboratory (stock 011122 on mixed genetic background). They were backcrossed on B6 background for more than 10 generations. All mice were 8-10 weeks old at the time of exposure. Animal breeding and study procedures were performed under an approved institutional animal care and use committee (IACUC) protocol at Duke University (A053-21-03). All animal experiments were conducted in accordance with the American Association for the Accreditation of Laboratory Animal Care guidelines.

### Rodent Exposures

Filtered air (FA) and ozone (O_3_) exposures followed published protocols.^21–24^ Exposures to FA or O_3_ (2 parts per million) for 3h were performed in 55-liter Hinners-style exposure chambers. The mice were exposed in stainless steel wire mesh cages inside the exposure chamber with dividers to prevent congregating. Chamber air was maintained at 20–22°C with 40–60% relative humidity and supplied at a rate of 20 changes/hour. O_3_ was generated by directing 100% oxygen through ultraviolet light. O_3_ concentration was continuously monitored by an O_3_ ultraviolet light photometer (Teledyne Instruments, model 400E). After 3h, the mice were removed from the chamber. For the lineage labeling experiments, Cx3cr1^ERcre^ zsGreen mice were administered 2 mg of tamoxifen via oral gavage. 24h after tamoxifen induction, the mice were exposed to FA or O_3_ and lungs were harvested 12, 24, or 72h post-exposure for flow cytometry to define lineage reporter expression. For clodronate studies, C57Bl/6J mice were dosed with 50uL of 5mg/mL clodronate-loaded liposomes (batch No. C12J0321 & C18E0422) (Liposoma BV) or PBS by oropharyngeal aspiration. 72h after clodronate or PBS, mice were exposed to FA or O_3_ and sacrificed 12 or 24h post-exposure. MerTK^-/-^ mice and control C57Bl/6J mice were exposed to FA or O_3_ (2 ppm x 3h) and harvested 24h post-exposure.

### Statistics

Statistical testing is as noted in the figure legends. In most settings, an ANOVA with post-hoc multiple comparisons analysis was conducted in PRISM 9 (GraphPad, Boston, MA) using Tukey’s Honestly Significant Difference Test to reveal individual comparisons within groups on both cell types and cytokines. A p-value of less than 0.05 was considered statistically significant. Data in figures are presented as individual data points ± standard error of the mean. To account for any variability within experiments and samples, we performed an additional secondary analysis. This used mixed effects linear regressions performed in MATLAB 2017b (MathWorks, Natick, MA). We assessed the effects of timepoint (12, 24h), exposure (FA, O_3_ (2 ppm x 3h)), and treatment (CL, PBS), and all 2-and 3-way interaction effects, on cell differentials, individual cytokines, and neutrophil elastase. The Robust Regression was fit using a bisquare weighting function that assigns observations with larger residuals a weight trending towards zero. The analysis conducted in MATLAB confirmed and validated the statistics run in PRISM.

### Additional methods can be found in the supplement

## Results

### Monocyte-derived AMØs are not recruited following acute O_3_ exposure in rodents

To define AMØ composition following acute O_3_ exposure in rodents, we utilized an established inducible monocyte/macrophage lineage tracing system (Cx3cr1^ERcre^ crossed to zsGreen mice).^17,18^ In this system, tamoxifen administration permanently labels nearly 100% of circulating monocytes, which express *Cx3cr1*, and their progeny (i.e. monocyte-derived AMØs), with a green fluorescent protein (GFP).^17^ In contrast, in Cx3cr1^ERcre^ zsGreen mice, tissue-resident AMØ, which do not express *Cx3cr1* in adulthood, are not labeled. Therefore, the Cx3cr1^ERcre^ zsGreen mice allow clear segregation of tissue-resident from monocyte-derived AMØs. To induce lineage tracing, tamoxifen was administered via oral gavage 24h prior to FA or O_3_ exposure (Figure 1A). The mice were harvested at 12, 24, and 72h post-exposure based on our and others’ prior studies defining peak and resolving time points of O_3_-induced inflammation and injury.^22,25,26^ Whole lung tissue was harvested and digested to obtain a single cell suspension and the cells were processed for flow cytometry using a previously published protocol (See supplemental Table 1 and 2 for antibodies and cytometer configuration).^22^ Lung MØs were defined as live, CD45^+^, Ly6G^-^, CD64^+^, CD24^-^ cells (Supplemental Figure 1). Based on CD11c versus CD11b expression, AMØs (CD11c^+^, CD11b^-^) were delineated from IMØs (CD11c^+/-^, CD11b^+^). AMØ definition was further confirmed using CD169 or CD206 (data not shown) and Siglec-F versus Ly6C (Supplemental Figure 1, blue panel). AMØs were then segregated by GFP expression to define tissue-resident AMØs (live, CD11c^+^, CD11b^-^, GFP^-^) versus monocyte-derived AMØs (live, CD11c^dim/-^, CD11b^+^, GFP^+^). Comparing GFP^+^ and GFP^-^ expression of AMØ in O_3_-exposed mice (Figure 1B and 1C), we observed that the majority of AMØs were GFP^-^, suggesting a predominance of tissue-resident AMØs and a lack of flux of monocyte-derived AMØs following O_3_ exposure. The red boxes depict the average percentage of monocyte-derived AMØ present in the FA (mean 1.154% +/-0.1724, n=3), 12h (mean 0.6352% +/-0.0765, n=5), 24h (mean 0.7240% +/-0.1201, n=4), and 72h (mean 0.1068% +/-0.1341, n=6) post-O_3_ exposure (Figure 1C). Despite the lack of monocyte-derived AMØ recruitment (Figure 1B), we did observe influx of monocytes following O_3_ exposure suggesting that monocytes traffic into the lung following exposure but do not differentiate into AMØs (Supplemental Figure 2). Overall, the data collected with an inducible monocyte lineage tracing mice show no evidence for lung recruitment of monocyte-derived AMØs following O_3_ exposure.

**Figure 1.**
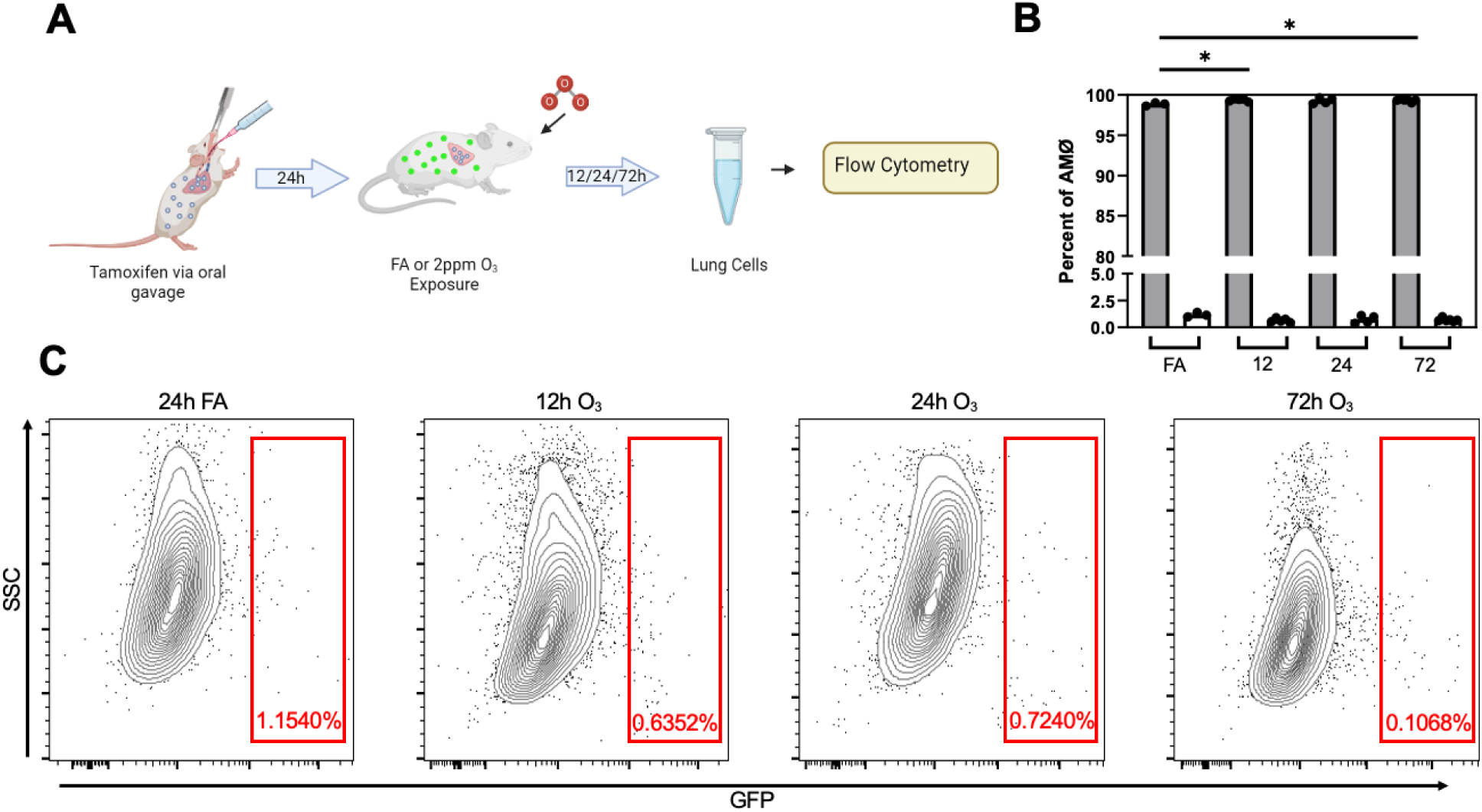
Monocyte-derived alveolar macrophages (AMØ) are not recruited following acute O_3_ exposure. **A**. Cx3cr1^ERCre^ x zsGreen mouse model was used. Circulating blood monocytes and monocyte-derived AMØ express Cx3cr1, and after tamoxifen induction are GFP^+^, while tissue-resident AMØ remain unlabeled. Lineage label was induced with tamoxifen 24h prior to exposure with filtered air or O_3_ (2 ppm) for 3h. 12, 24, or 72h-post exposure, whole lungs were processed and analyzed by flow cytometry. Schematic made using BioRender.com **B**. Bar plots represent individual mouse GFP^-^ (tissue-resident AMØ, grey) and GFP^+^ (monocyte-derived AMØ, white) populations. ANOVA analysis was conducted using Tukey’s Honestly Significant Difference Test post-hoc. Data points represented with SEM; showing significant increase of tissue-resident AMØ from FA to 12 and 72h post O_3_ exposure. *p<0.05. **C**. Acute O_3_ exposure does not induce recruitment/differentiation of monocyte-derived AMØ. Representative flow cytometry plots of AMØs were negative for zsGreen/GFP positive cells, supporting no evidence of monocyte-derived AMØ recruitment. The red boxes indicate where GFP^+^ cells (monocyte-derived AMØ) would be observed if they were present in these samples. Percent GFP^+^ indicated within the respective box in red. Full flow cytometry gating strategy and antibodies can be found in the supplements (Supplemental Figure 1). Flow plots are representative of n=3-6 mice per timepoint and replicated x1. Mice used include 4 females at 12h, with all other mice being male.

### Monocyte-derived AMØs are not recruited following acute O_3_ exposure in healthy human volunteers

Given the lack of monocyte-derived AMØ recruitment following acute O_3_ exposure in rodents, we wanted to translate this observation to humans undergoing acute laboratory O_3_ exposure. Healthy human volunteers (n=12) without a history of chronic cardiopulmonary disease (Figure 2A) were exposed to FA or O_3_ (200 ppb) for 135 minutes with intermittent exercise in a randomized crossover study design with an 18–20 day washout period between exposures. Approximately 21h following each exposure, the subjects underwent a bronchoscopy with bronchoalveolar lavage (BAL). 10mL of BAL fluid was aliquoted per exposure and processed/stained/analyzed by flow cytometry to define AMØ composition. Human AMØs were defined as singlets, live, CD45^+^, CD3^-^, CD15^-^, CD206^+^ cells (Figure 2B and Supplemental Table 3). Segregation of human tissue-resident from monocyte-derived AMØs was defined by the extent of CD206 and CD14 staining where tissue-resident AMØs are CD206^Hi^, CD14^Lo^ and monocyte-derived AMØs are CD206^Lo^, CD14^Hi^ (Figure 2B). Monocytes were segregated and identified as CD14^Hi^, finding no significant monocyte recruitment following O_3_ exposure (Figure 2B and 2C). Consistent with prior human data,^27^ healthy human volunteers had monocyte-derived AMØs in the FA exposure samples (Figure 2C). However, following O_3_ exposure there was no significant recruitment of monocyte-derived AMØs (Figure 2B and 2C). Despite the lack of overall evidence for recruitment of monocytes or monocyte-derived AMØs, we did observe significant intra-individual variability in filtered air and O_3_ exposure immune cell profiles, suggesting differences in basal airspace immune cell composition and response to exposure between study subjects (Figure 2D). Overall, this data suggests that similar to mice, healthy human volunteers do not recruit monocyte-derived AMØs following acute laboratory O_3_ exposure.

**Figure 2.**
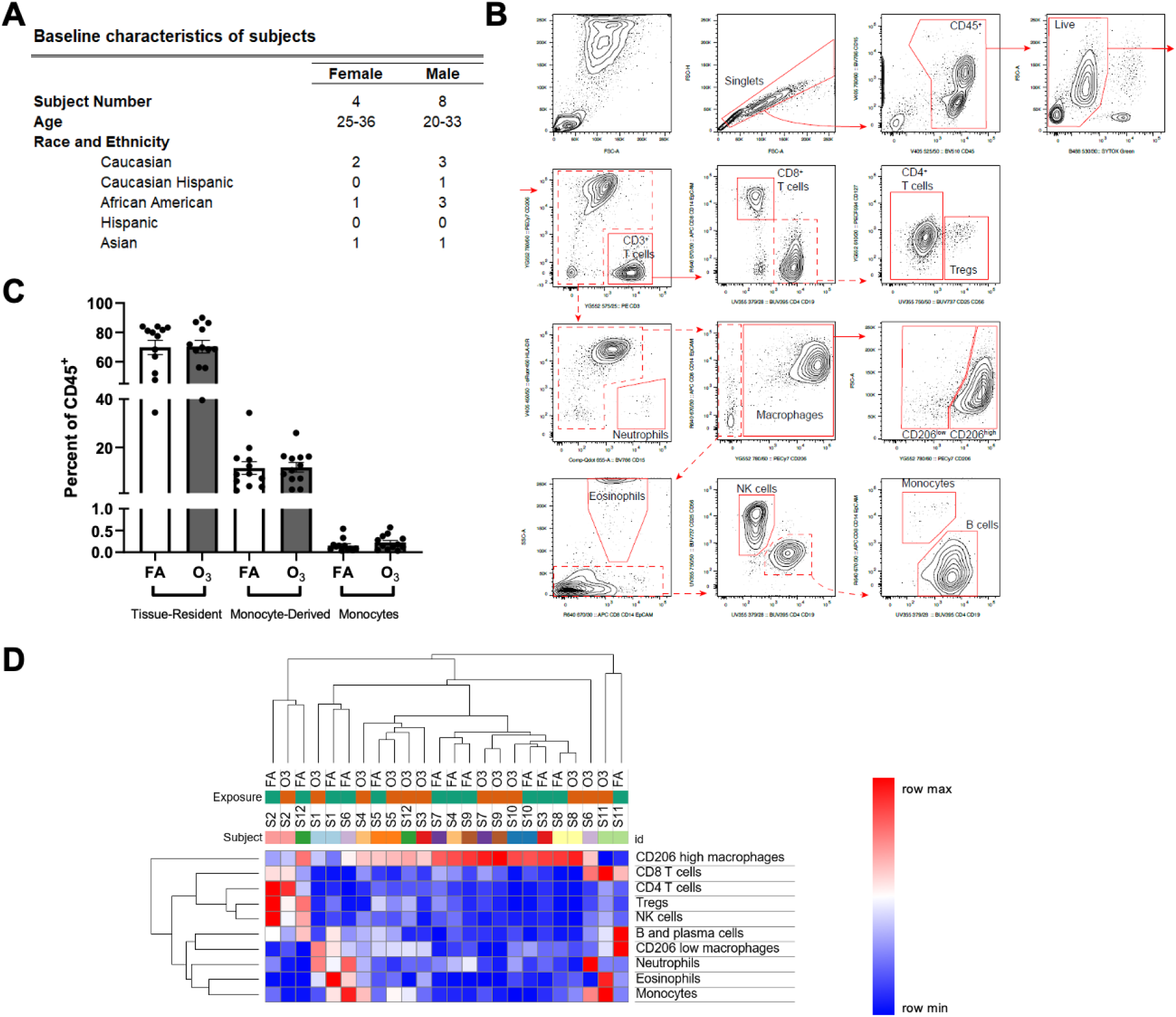
Monocyte-derived alveolar macrophages (AMØ) are not recruited following acute O_3_ exposure in humans. A. Human subjects were exposed to FA and O_3_ (2ppm) for 3h in separate visits. They then underwent a bronchoscopy in which BAL fluid was collected and processed and analyzed by flow cytometry. Demographic information is listed for all human subjects including sex, age, and ethnicity. B. Representative flow cytometry plots of the gating strategy used to identify alveolar macrophages were generated demonstrating populations of CD206^Lo^, monocyte-derived AMØ, and CD206^Hi^, tissue-resident AMØ, populations. The CD206^Lo^ population did not increase following acute O_3_ exposure, supporting no evidence of monocyte-derived AMØ function in this context. The red boxes indicate where GFP+ monocyte-derived AMØ would be observed if they were present in these samples C. Acute O_3_ exposure does not induce recruitment of monocyte-derived AMØ. Bar plots represent human cellular response to FA (white) and O_3_ (grey) populations. Paired t-test analysis was conducted and data points are represented with SEM. No significant difference seen in tissue-resident AMØ, monocyte-derived AMØ, and monocyte populations post O_3_ exposure. **D**. Hierarchical clustering of the flow cytometry data from BAL samples. Column headers are color-coded by the exposure type (FA or O_3_) and subject. Samples were clustered by Euclidean distance using average linkage method. *p<0.05 Flow plots are representative of n=12 human subjects.

### Tissue-resident AMØ depletion causes persistent O_3_-induced bronchoalveolar lavage neutrophilia

Given the lack of monocyte-derived AMØ recruitment following O_3_ exposure in rodents or humans, we focused on defining a role of tissue-resident AMØs in acute O_3_ exposure. Following previously published protocols,^16^ tissue-resident AMØs were depleted using administration of intra-tracheal clodronate-loaded liposomes. To confirm specificity of this clodronate-mediated depletion, we performed whole lung flow cytometry assessment of immune cells, identifying that clodronate-loaded liposomes led to AMØ-specific depletion (Figure 3A). Administering clodronate-loaded liposomes to tamoxifen-induced Cx3cr1^ERcre^ zsGreen mice, we confirmed that clodronate administration did not cause recruitment of monocyte-derived AMØs during the timescale of our exposure experiments (Figure 3B). The red boxes depict the average percentage of monocyte-derived AMØ present (mean 0.9200% +/-0.3107, n=4). This data demonstrates that clodronate-mediated depletion was specific to AMØs and resulted in depletion of tissue-resident AMØs without subsequent recruitment of monocyte-derived AMØs.

**Figure 3.**
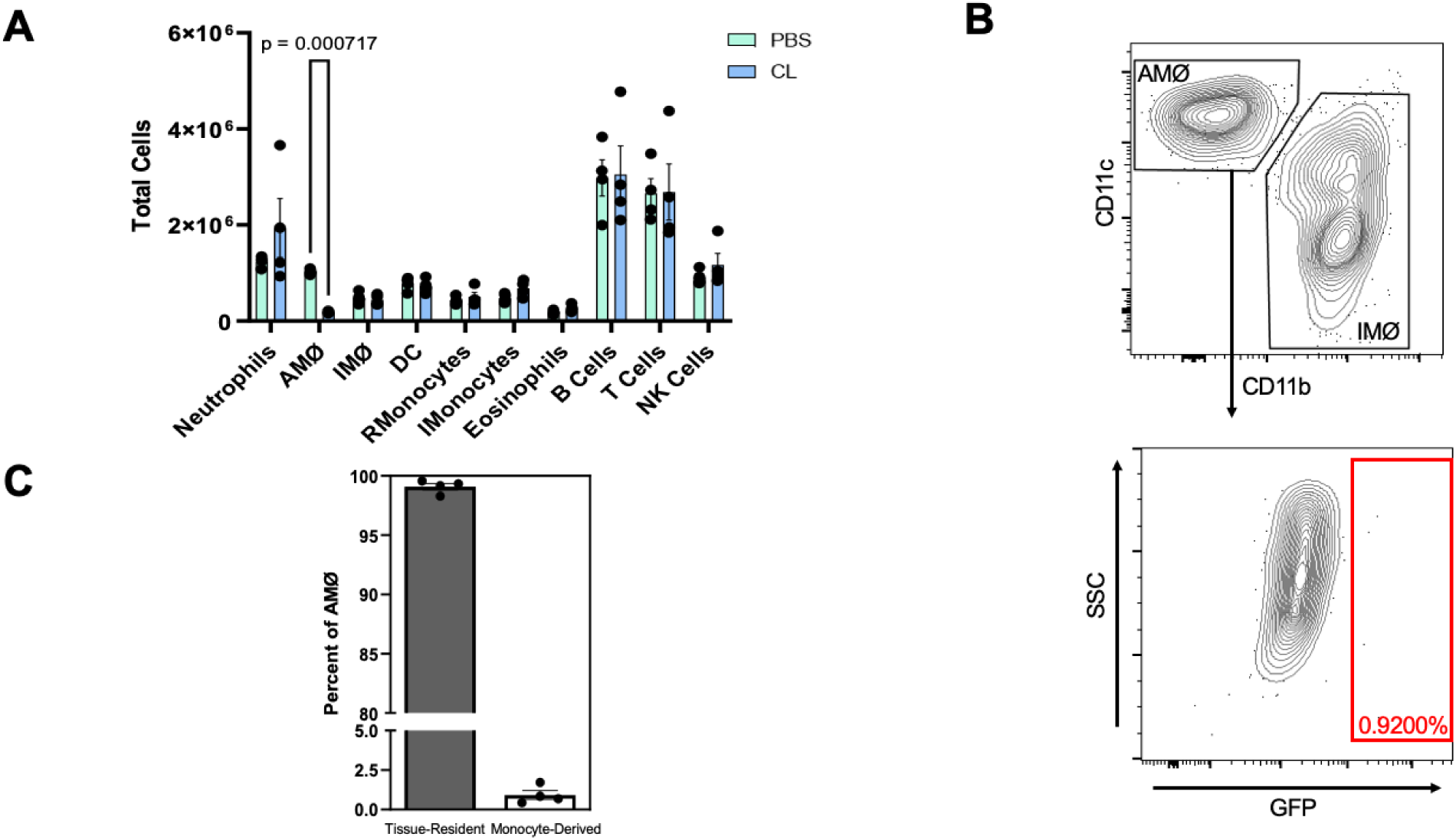
Airspace administration of clodronate (CL) depletes tissue-resident alveolar macrophages (AMØ). A. C57Bl/6J female mice were dosed with clodronate or PBS (control) were harvested 72h post-dosing. The whole lung tissue samples were perfused, digested, and stained to identify the individual cells via flow cytometry. Bars shaded green indicate PBS/vehicle control while blue indicates CL administration. Data is from n=4 mice per group, p value is as identified. **B.** Cx3cr1^ERCre^ x zsGreen mice were dosed with CL 72h before harvest then underwent lineage labeling with tamoxifen 48h post-dosing. They were then harvested and processed for flow cytometry. The lack of GPF^+^ AMØ (red box) suggests a lack of Cx3cr1^ERCre^ x GFP^+^ monocyte-derived AMØs as a result of CL depletion of tissue-resident AMØ. Percent GFP^+^ indicated within the respective box in red. Flow plot is a representative sample of n=4. **C.** Bar plot represents individual mouse GFP^-^ (tissue-resident AMØ, grey) and GFP^+^ (monocyte-derived AMØ, white) populations, indicating no recruitment of monocyte-derived AMØ following CL depletion. Full flow cytometry gating strategy and antibodies can be found in the supplements (Supplemental Figure 1).

Next, we assessed the impact of clodronate-mediated depletion of tissue-resident AMØs on acute O_3_-induced lung inflammation and injury. C57BL/6J mice were administered PBS or clodronate and then 72h post-administration were exposed to FA or O_3_ (2 ppm) for 3h (Figure 4A). At 12h or 24h post-exposure, mice were harvested and assessed for airspace inflammation. No differences in total cells, macrophages, or neutrophils were observed in the filtered air groups (Figure 4B, FA). In PBS-administered, O_3_-exposed mice, there was an increase in BAL total cells at both 12 and 24h post-exposure (Figure 4B). Cell differentials revealed an increase in airspace neutrophils that peaked at 12h and had started to decline by 24h post-exposure. In contrast, in the clodronate-administered, O_3_-exposed mice, neutrophils increased at 12h post-exposure but then remained persistently elevated at 24h post-exposure (Figure 4B). As expected, AMØs were reduced at 12h and remained so through 24h post-exposure in mice given clodronate administration prior to O_3_. In contrast, there was a significant difference in the amount of airspace neutrophils 24h post-exposure in the PBS and clodronate administered mice exposed to O_3_, whereas the clodronate administered mice demonstrated persistent airspace neutrophilia.

**Figure 4.**
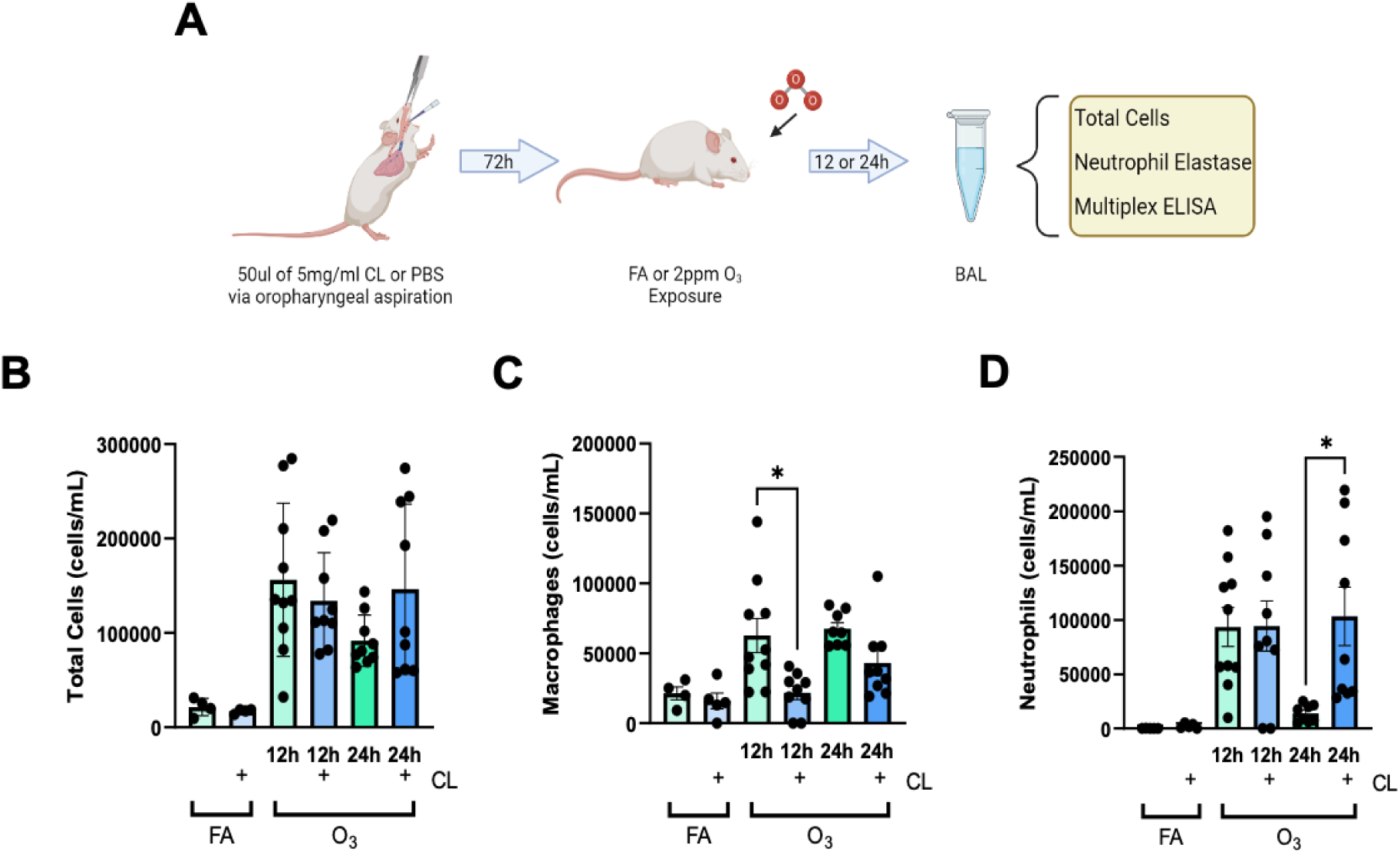
Tissue-Resident AMØ depletion leads to persistent O_3_-induced BAL neutrophilia. **A.** Male mice were dosed with 50uL of 5mg/ml clodronate liposomes via oropharyngeal aspiration. 72h following clodronate depletion of tissue-resident AMØ, mice were exposed to FA or O_3_ (2 ppm) for 3h. Mice were then harvested 12/24h following exposure and the samples were processed for cell differentials and other measures of inflammation. Schematic made with BioRender.com. **B.** BAL total cells, (**C.**) macrophages, and (**D.**) neutrophils were enumerated following PBS or Clodronate (CL) administration and FA or O_3_ exposure. Bars shaded green indicate PBS/vehicle control while blue indicates CL administration. By total cell numbers, CL exposed mice exhibited a reduction of macrophages at 12h and increased neutrophils at 24h. n=4-10 mice per group/exposure/timepoint *p<0.05. ANOVA analysis was conducted using Tukey’s Honestly Significant Difference Test post-hoc.

### Tissue-resident AMØ depletion causes persistence of BAL cytokine/chemokines and neutrophil elastase

In addition to airspace inflammation, we also assessed the impact of tissue-resident AMØ depletion on other measures of O_3_-induced lung inflammation and injury. BAL fluid cytokines and chemokines levels were measured by multiplex ELISA with a focus on neutrophil factors. These included Granulocyte colony-stimulating factor (G-CSF), Leukemia inhibitory factor (LIF), IP-10 (also known as CXCL10), interleukin (IL)-6, GROa (also known as CXCL1), Monocyte chemoattractant protein-1 (MCP-1 or CCL2), Macrophage inflammatory protein 2-alpha (MIP-2α), and Macrophage inflammatory protein-2 beta (MIP-2β). At 12h post-exposure, LIF, IL-6, GROa, MIP-2α, and MIP-2β were increased when compared to FA groups but no difference was observed between PBS or clodronate administered groups. Alternatively, at 24h post-O_3_ exposure, differences were present in the cytokine responses between PBS and clodronate administered groups (Figure 5), where IP-10, IL-6, GROa, MCP-1, MIP-2α, and MIP-2β were all elevated in clodronate administered but not control mice. Similar to cytokine responses, BAL neutrophil elastase was elevated in O_3_ exposed mice at 12h when compared to FA but not different based on clodronate administration (Supplemental Figure 3). However, neutrophil elastase was persistently increased at the 24h post-exposure time point in the clodronate administered versus control mice. This data supports that in tissue-resident AMØ-depleted mice, the O_3_ exposure induced persistence in airspace neutrophils associates with a similar persistence in pro-inflammatory cytokines and neutrophil elastase suggesting failed resolution of O_3_-induced inflammation.

**Figure 5.**
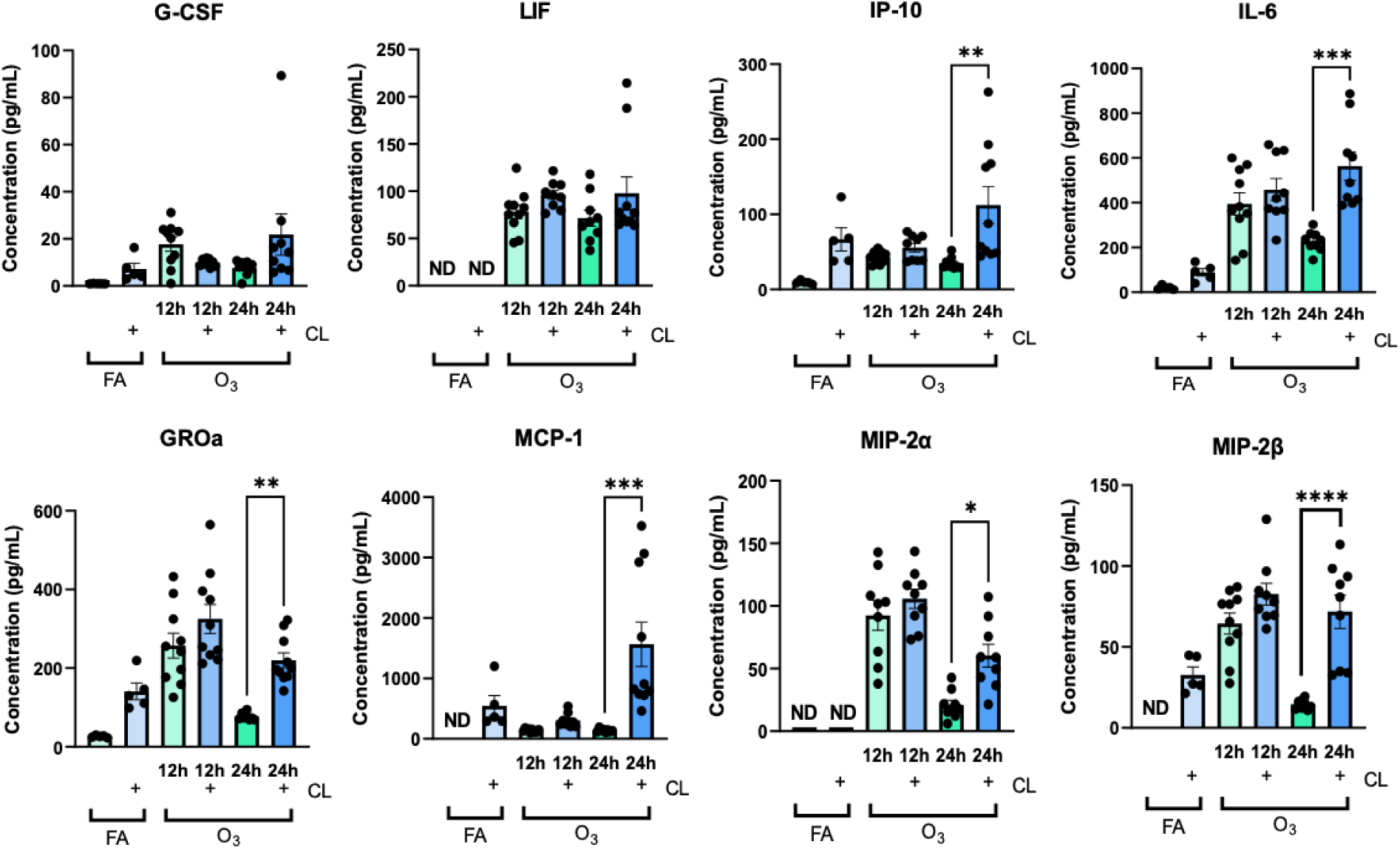
Increased BAL cytokines associated with neutrophil recruitment in O_3_-exposed tissue-resident AMØ depleted mice. BAL fluid from PBS or Clodronate and FA or O_3_ (2ppm) exposed male mice was assessed for cytokine expression by multiplex ELISA for inflammatory cytokines and known neutrophilic chemotactic factors. Bars shaded green indicate PBS/vehicle control while blue indicates CL administration. n=4-10 mice per group/exposure/timepoint *p<0.05. ANOVA analysis was conducted using Tukey’s Honestly Significant Difference Test post-hoc. ND indicates samples below the assay limit of detection.

### Depletion of tissue-resident AMØs reduces efferocytosis in vivo

The persistence of airspace neutrophils following O_3_ exposure in tissue-resident AMØ-depleted mice suggests failed clearance of apoptotic neutrophils. To determine if neutrophils exhibit increased apoptosis following clodronate-loaded liposome administration, we assessed apoptosis markers (7AAD and Annexin V) by flow cytometry in Ly6G^+^ BAL neutrophils. We observed that O_3_ exposed PBS administered control mice had a higher portion of “Live” neutrophils (Ly6G^+^, 7AAD^-^, Annexin V^-^) and reduced “Early Apoptosis” neutrophils (Ly6G^+^, 7AAD^-^, Annexin V^+^) than O_3_ exposed clodronate-loaded liposome administered mice (Supplemental Figure 4). This finding suggests that depletion of tissue-resident AMØs resulted in increased apoptotic neutrophils following O_3_ exposure, supporting defective efferocytosis. To confirm the requirement of tissue-resident AMØs for *in vivo* efferocytosis, we adapted a method from Hodge et al. to assess efferocytotic function *in vivo*.^28^ Apoptotic cells were generated by UV irradiating immortalized T cells (Jurkat cells) and were labeled with Calcein AM dye to track their clearance. Prior to dosing, the extent of Jurkat cell apoptosis was confirmed by viability staining (data not shown). PBS or clodronate was administered to C57Bl/6J mice as described above. 72h after administration, apoptotic labeled Jurkat cells, were instilled via oropharyngeal aspiration and 1.5h later mice were harvested for BAL allowing for the number of labeled Jurkat cells to be measured by flow cytometry (Figure 6A). To assess *in vivo* efferocytosis we created a ratio of the number of recovered labeled Jurkat cells in the BAL over the number of instilled Jurkat cells. We observed that the mice with tissue-resident AMØ depletion exhibited decreased efferocytosis, as evidenced by an increased number of Jurkat cells recovered from BAL fluid (Figure 6B). This supports that tissue-resident AMØ facilitate effective efferocytosis.

**Figure 6.**
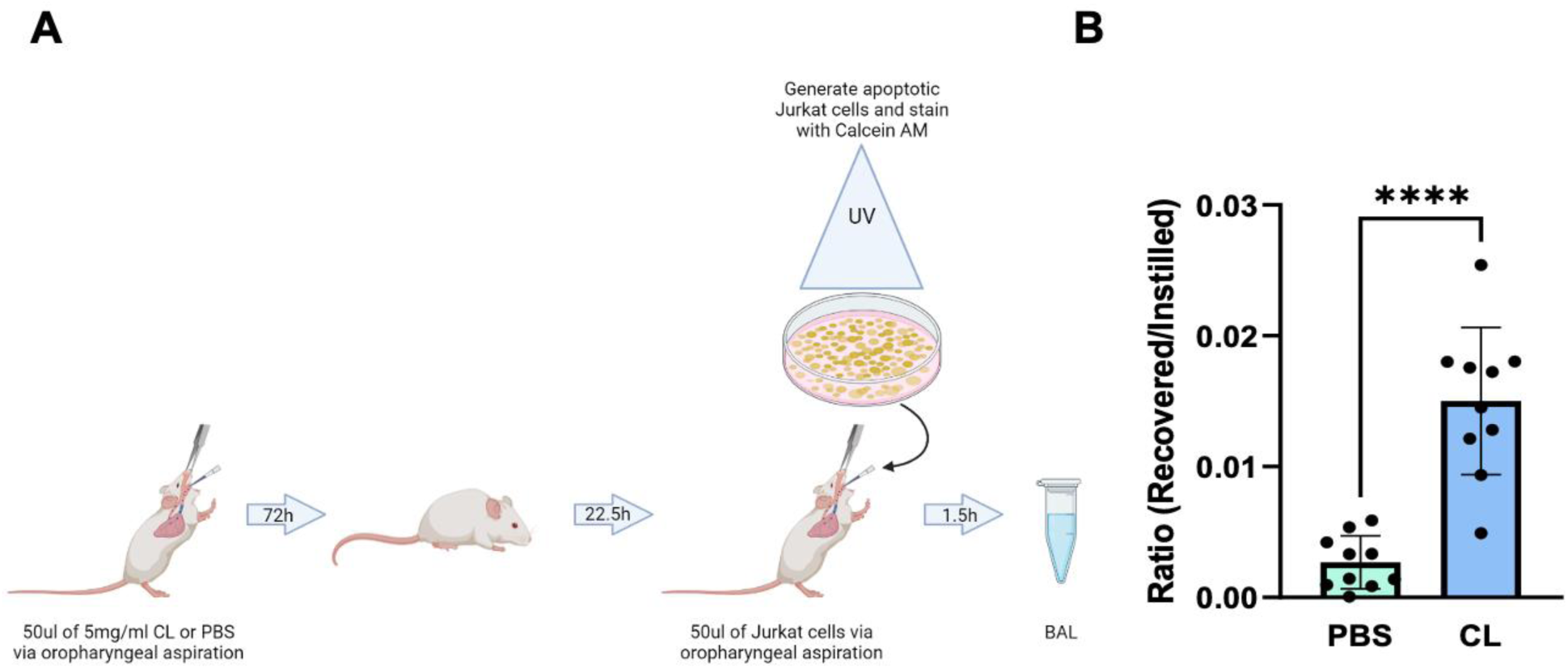
Tissue-resident alveolar macrophage (AMØ) depletion decreased efferocytosis. **A.** C57BL/6J male mice were dosed with CL or PBS. The mice were then instilled with apoptotic immortalized T cells, Jurkat cells, via oropharyngeal aspiration 22.5h after exposure for 1.5h immediately prior to the harvest. During the harvest, 24h post-exposure, BAL was collected and prepared for use in flow cytometry. Schematic made with BioRender.com. **B.** Apoptotic cell clearance was defined using Calcein AM, positive cells indicating live or early apoptotic cells. Healthy cells would not be engulfed by AMØs, thus decreased Calcein AM positive cells indicate clearance of apoptotic cells (efferocytosis). The graphs depict the ratio of counted recovered vs. instilled cells from the collected BAL. Bars shaded green indicate PBS/vehicle control while blue indicates CL administration. An unpaired T-test was conducted with n=10 mice per group/exposure/timepoint *p<0.05.

### Mer tyrosine kinase (MerTK) null mice exhibit persistent O_3_-induced inflammation

As tissue-resident AMØs depleted mice exhibited decreased efferocytosis, we wanted to validate the role of efferocytosis in acute O_3_ exposure responses. To perform this, we assessed O_3_ exposure responses in MerTK null (MerTK^-/-^) mice. MerTK cell surface staining is a defining feature of AMØs and is frequently used in flow cytometry studies to segregate AMØs from other lung immune cells.^29^ In addition, MerTK regulates efferocytosis and MerTK^-/-^ mice are noted to have ineffective efferocytosis.^19,20^ We first assessed if MerTK expression was modified by O_3_ exposure. We measured MerTK expression by real time PCR from BAL cells at 6h, 12h, 24h, 48h, 72h and 7 days post-acute O_3_ exposure (2 ppm for 3h). Overall, there was minimal overall change in MerTK expression (Figure 7A). We then compared O_3_-induced lung inflammation and injury responses in C57Bl/6J (WT) and MerTK^-/-^ mice. WT and MerTK^-/-^ mice were exposed to FA or O_3_ (2 ppm) for 3h and then harvested 24h post-exposure. We focused on the 24h time point since this is where we identified differences in the resolution of O_3_-induced neutrophilic inflammation. No differences were observed in BAL immune cells in FA exposed WT and MerTK^-/-^ mice. Following O_3_ exposure, MerTK^-/-^ mice, compared to WT mice, demonstrated increased total cells, macrophages, and neutrophils (Figure 7B). Furthermore, BAL cytokine and chemokine profiles were enhanced in the O_3_ exposed MerTK^-/-^ mice when compared to WT mice (Figure 8). This included elevations in IP-10, IL-6, GROa and MIP-2β. This suggests that MerTK promotes resolution of O_3_-induced inflammation.

**Figure 7.**
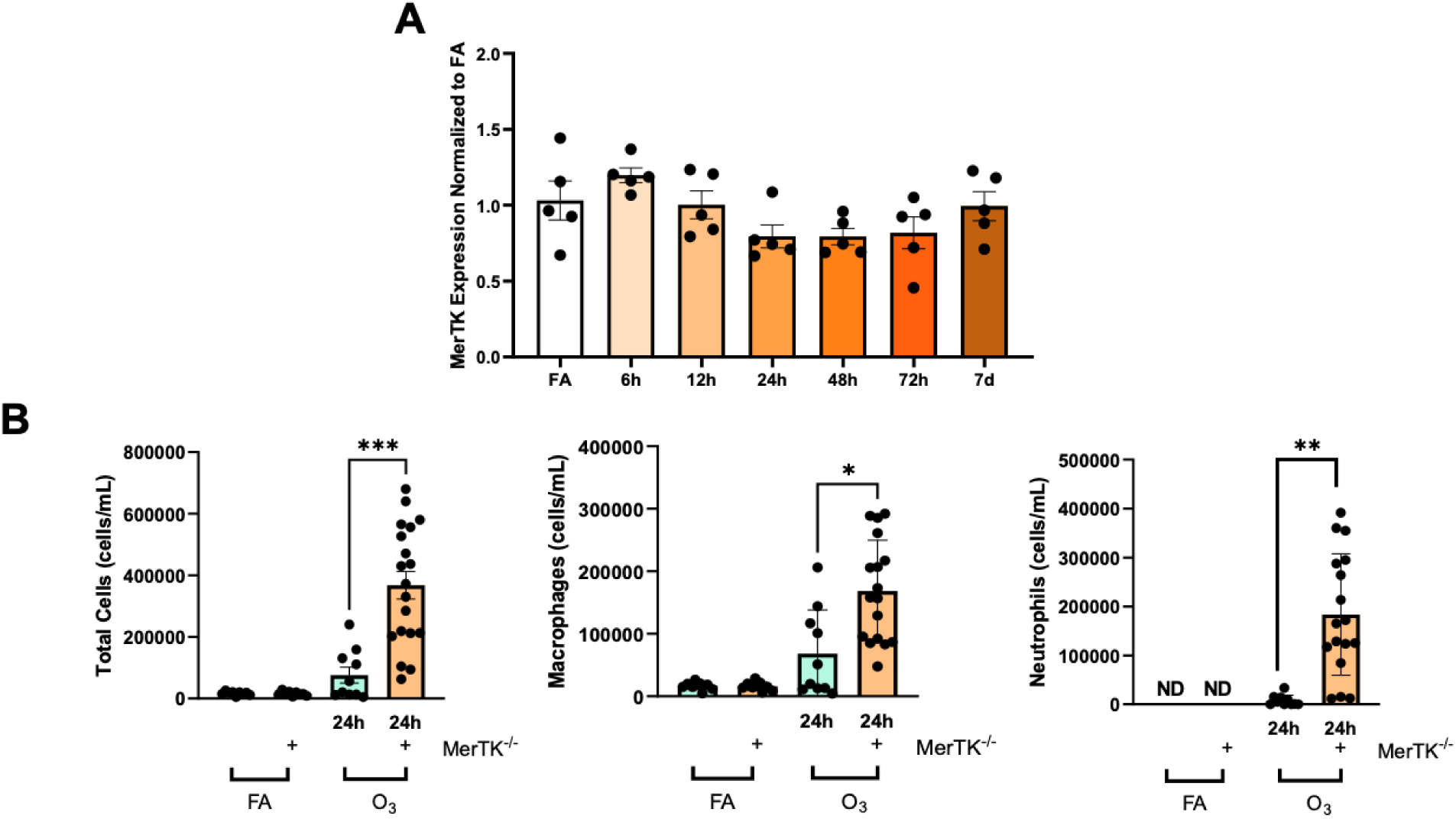
MerTK^-/-^ mice have increased O_3_-induced airspace inflammation. A. C57BL/6J male mice were exposed to O_3_ (2 ppm) for 3h and harvested 6h, 12h, 24h, 48h, 72h, and 7days later. mRNA expression of MerTK was assessed using the collected BAL cells. n=5 mice per time point. **B.** MerTK^-/-^ mice and C57BL/6J mice (wildtype) were exposed to FA or O_3_ (2 ppm) for 3h. The mice were harvested 24h post-exposure and the BAL was collected and processed for total cell counts and differentials. Green bars indicate the C57BL/6J mice while the orange indicates the MerTK^-/-^ genotype. n=9-19 mice per group. *p<0.05. ANOVA analysis was conducted using Tukey’s Honestly Significant Difference Test post-hoc. ND indicates samples below the limit of assay detection.

**Figure 8.**
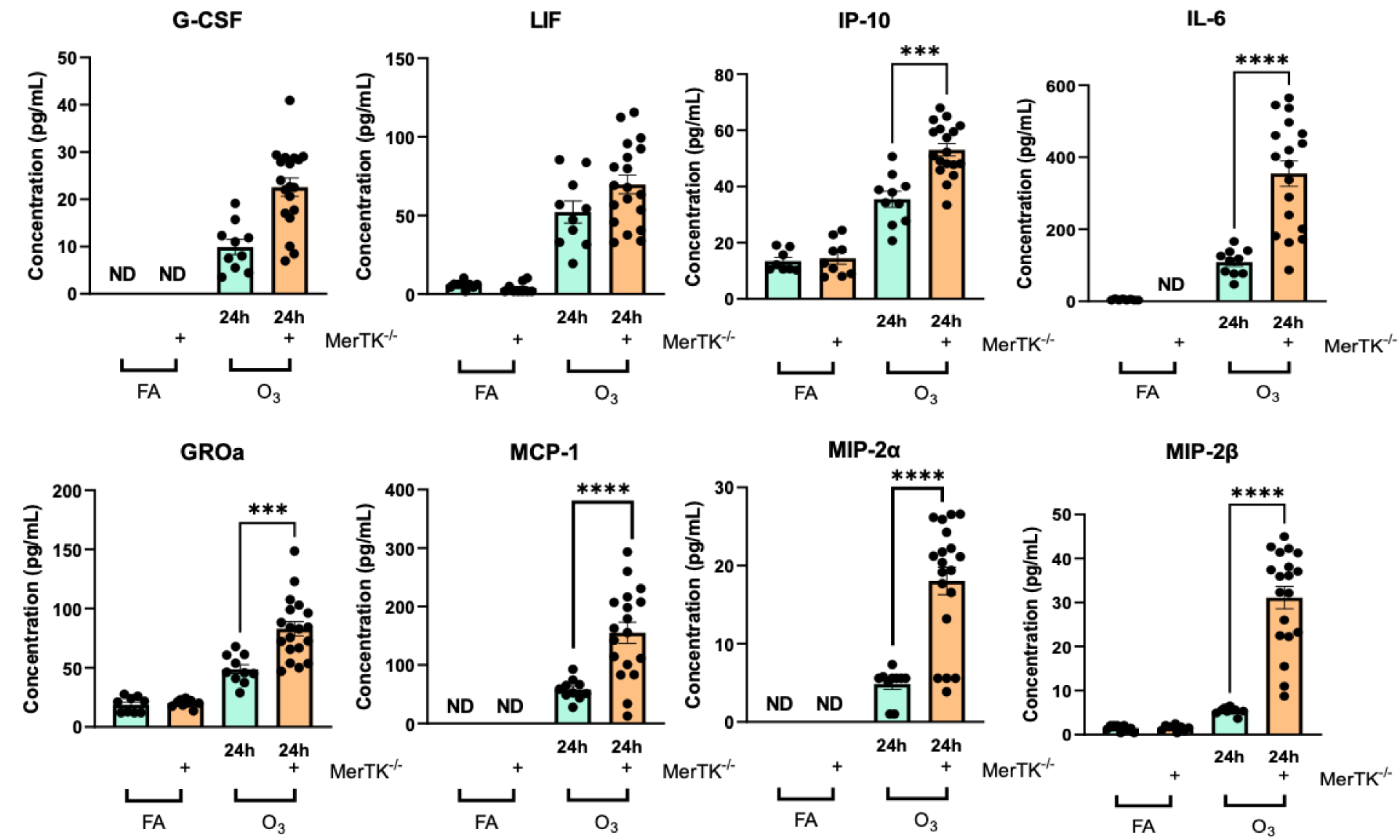
Increased BAL cytokines associated with neutrophil recruitment in O_3_-exposed MerTK^-/-^ mice. BAL fluid from C57BL/6J and MerTK^-/-^ mice FA or O_3_ (2ppm) exposed mice was assessed for cytokine expression by multiplex ELISA for inflammatory cytokines and known neutrophilic chemotactic factors. Green bars indicate the C57BL/6J mice while the orange indicates the MerTK^-/-^ genotype. n=8-19 mice per group/exposure/timepoint. *p<0.05. ANOVA analysis was conducted using Tukey’s Honestly Significant Difference Test post-hoc. ND indicates samples below the limit of assay detection.

## Discussion

AMØs are critical to the initiation, propagation, and resolution of lung inflammation and injury. Therefore, defining specific AMØ functions in diverse lung injury responses remains an important research question. In the present study, we focused on defining the role of AMØs in O_3_-induced lung injury. We performed detailed lineage tracing of AMØs following O_3_ exposure to define the origin of AMØs. We identified that following O_3_ exposure AMØs are of tissue-resident origin without a predominance of monocyte-derived AMØs. Similar to observations in rodents, healthy humans exposed to O_3_ also did not have an increase in monocyte-derived AMØs. To define the function of tissue-resident AMØs, we performed a series of experiments using a clodronate-mediated tissue-resident AMØ depletion model to demonstrate that tissue-resident AMØs facilitate efferocytosis following O_3_ exposure and that efferocytosis promotes resolution of O_3_-induced lung injury. To confirm the requirement of effective efferocytosis in the resolution of O_3_-induced lung inflammation, we performed O_3_ exposure studies in MerTK^-/-^ mice, which have a genetic deficiency in efferocytosis. Following O_3_ exposure, MerTK^-/-^ mice exhibited persistent airspace neutrophils and cytokine responses similar to exposed tissue-resident AMØ depleted mice. Overall, the data support that O_3_ exposure does not recruit monocyte-derived AMØs and tissue-resident AMØs appear to promote resolution of O_3_-induced lung injury through efferocytosis via MerTK.

Acute O_3_-induced lung injury is regulated by AMØ functions, with data supporting both pro-inflammatory and protective/pro-resolving functions.^3,30–32^ While, AMØ origin may offer one potential explanation for differing functions of AMØs following O_3_ exposure, there have been inconsistent findings about AMØ composition following O_3_ exposure. Francis et al. defined populations of pro-inflammatory (CD11b^+^ Ly6C^Hi^ and iNOS^+^) and anti-inflammatory (CD11b^+^ Ly6C^Lo^) AMØs present in the lung following O_3_ exposure, which required CCR2 for their recruitment.^33^ In addition, another study by this group identified that these O_3_-induced pro-inflammatory AMØs were reduced in splenectomized mice suggesting these pro-inflammatory AMØs are derived from a splenic intermediates.^34^ Overall, their data on pro-inflammatory AMØs suggests these cells are monocyte-derived in origin (i.e. monocyte-derived AMØ) due to their CCR2 requirement and origin from a splenic intermediate. As CCR2-null or splenectomized mice exhibited reduced O_3_-induced inflammation and injury, the authors argued that these pro-inflammatory AMØ were driving O_3_-induced pathobiology. Alternatively, our prior work suggests that O_3_-derived AMØs are principally tissue-resident in origin and exhibit a protective role in O_3_-induced lung injury.^23^ We defined expansion of a Cx3cr1-mediated tissue-resident AMØ subset following O_3_ and loss of this subset in Cx3cr1-null mice exacerbated O_3_-induced lung inflammation and airway hyperresponsiveness.^23^ In a separate study, we compared O_3_ exposure responses in male and female mice using a more extensive immune phenotyping flow cytometry panel and also did not identify monocyte-derived AMØs.^22^ However, none of these studies relied on lineage tracing strategies that clearly define AMØ origin. To address this, the present study used lineage tracing with inducible Cx3cr1^ERcre^ zsGreen mice, which distinguishes tissue-resident AMØ from monocyte-derived AMØs.^17^ We identified that AMØs following O_3_ exposure are predominantly tissue-resident in origin with minimal to no monocyte-derived AMØs (Figure 1C). This supports that macrophage-derived effects in lung O_3_ responses are restricted to tissue-resident AMØs.

Though we did not observe an influx of monocyte-derived AMØs, we did observe influx of monocytes, but these monocytes did not differentiate into macrophages (Supplemental Figure 2). Prior work by Jakubzick et al. identified that monocytes can traffic through lung tissues without differentiation into macrophages and have antigen presentation functions.^35,36^ Presently, we do not know the specific functions of monocytes present following O_3_ exposure although it is possible that they could be driving some of the pro-inflammatory functions ascribed to macrophages following O_3_ exposure. Future studies will need to consider the functions of these monocytes in O_3_-induced lung inflammation and injury.

Due to the lack of monocyte-derived AMØ recruitment following acute rodent O_3_ exposure, we were interested to define if a similar AMØ responses would be observed in O_3_-exposed human subjects. Using BAL cells obtained by bronchoscopy from a cohort of healthy human volunteers exposed to FA and O_3_, we were able to confirm that monocyte-derived AMØs were not recruited (Figure 2C). Consistent with other healthy human BAL studies, we observed that monocyte-derived AMØ subsets are present in control human BALs.^27^ This is different from rodents, where AMØs have been shown to be of tissue-resident origin. This has been hypothesized to be a consequence of differences between laboratory rodents living in largely sterile housing environments and human volunteers living in non-sterile environments with exposures to pathogens and irritants.^37^ Understanding if these human monocyte-derived cells are hyper-responsive to subsequent exposures and/or pathogens remains an unanswered question and will need to be considered in future studies. One limitation of our flow cytometry analysis is that we did not define subsets of tissue-resident and/or monocyte-derived AMØs. This is despite single cell RNA-sequencing identifying 2 or more unique subsets distinct clusters of tissue-resident and monocyte-derived AMØs with the potential to have distinct functions.^27,38^ Future single cell RNA-sequencing studies will need to consider if there are distinct exposure related subsets of AMØs and their role in exposure human exposure responses.

Though we did not observe an overall difference in monocyte-derived AMØs following O_3_ exposure, evaluation of the immune cell profiles between individuals suggested significant diversity in immune cell profiles (Figure 2D). Comparing profiles across individuals in the FA controls, we observed significant differences in immune cell profiles, highlighting baseline intra-individual variability in airspace immune cell composition. Furthermore, we observed additional variability in FA to O_3_ exposure responses, where some individuals had minimal change and others clear immune profile differences between their FA and O_3_ exposures. This highlights intra-individual variability in baseline immune cell composition and in exposure responses. Future studies will need to consider if this baseline composition predicts the exposure responses or if exposure immune cell profiles predict individuals with adverse outcomes to exposure.

An interesting aspect of the present study was the observation that monocyte-derived AMØs were not recruited following acute O_3_-induced lung injury (Figure 1B and 1C). This is counter to other lung injury models where monocyte-derived AMØs predominate both in cell numbers and injury-regulating functions.^16,17^ This may reflect the relative severity of O_3_-induced lung injury versus other lung injury models. In C57Bl/6J mice, O_3_ exposure is typically associated with mild resolving lung injury.^31^ This is in contrast to other commonly used lung injury models such as LPS, bleomycin, and influenza infection, which are associated with extensive tissue injury and destruction.^13,39,40^ Generally, in these lung injury models, tissue-resident AMØs are largely dispensable, and principal injury-regulating effects have been ascribed to monocyte-derived AMØs. Although not specifically evaluated in the present study, it is possible that different mechanisms are driving responses in these more severe forms of lung injury versus mild and resolving lung injuries. This has important potential human health implications as mild lung injuries, such as those experienced following O_3_ exposure, are ubiquitous and therefore more commonly experienced than severe lung injuries. Understanding distinct mechanisms driving mild versus more severe lung injuries could shed light on how mild, normally resolving lung injury responses can become more severe and/or persistent. This insight may help us comprehend how ubiquitous injuries that typically resolve, like O_3_ exposure, can cause or exacerbate lung diseases. Consistent with this, specific genetic mouse strains exhibit enhanced sensitivity to O_3_ exposure, driving enhanced lung injury, and in some cases the development of lung fibrosis.^41,42^ Given this importance, future studies will need to consider additional mechanisms regulating mild lung injury that favor return to homeostasis and how these might be dysregulated under certain conditions to promote lung disease.

Given the lack of monocyte-derived AMØs, we focused on tissue-resident AMØ function in O_3_-induced acute lung injury. Using a combination of pharmacologic (i.e. clodronate loaded liposome-mediated tissue-resident AMØ depletion) and genetic (i.e. MerTK null mice) methods, we demonstrated a key role for tissue-resident AMØs in promoting resolution of O_3_-induced lung inflammation via efferocytosis. We identified that tissue-resident AMØ depletion prior to O_3_ exposure led to persistence of airspace neutrophilia (Figure 4B) and these neutrophils demonstrated increased apoptosis markers (Supplemental Figure 4) supporting defective apoptotic neutrophil clearance. We then directly demonstrated that tissue-resident AMØs are required for efferocytosis *in vivo* (Figure 6). To confirm the requirement for efferocytosis in resolution of acute O_3_-induced lung injury, we compared FA and O_3_ exposures in MerTK null mice. We observed that O_3_-exposed MerTK null mice, similar to O_3_-exposed clodronate-depleted tissue-resident AMØ mice, demonstrated persistent airspace neutrophilia and production of cytokines (Figure 8). Though, MerTK function has been studied in acute lung injury,^43,44^ to our understanding, this is the first description of MerTK function following acute O_3_-exposure. Interestingly, though we did not observe an overall change in BAL cell MerTK expression, there was a trend towards reduced MerTK at 24h post-exposure. The relative decrease in MerTK at 24h is the same time point where prior data has identified O_3_-induced reductions in efferocytosis.^28^ It is possible this reduction in efferocytosis could be a result of our observed changes in MerTK expression. Overall, this data suggests that tissue-resident AMØ promote the resolution of O_3_-induced lung inflammation via efferocytosis, and this occurs via a MerTK-mediated mechanism. Future studies could consider if augmentation of MerTK expression could improve O_3_-induced reductions in efferocytosis and if this could be a therapeutic intervention to promote resolution of O_3_-induced lung injury.

There are limitations to our study that should be considered. Despite the lack of clodronate-mediated impact on immune cell composition (Figure 2A), it is possible that clodronate might impact the function of non-AMØ immune cell subsets. Consistent with this, Culemann et al. recently identified that clodronate ingestion by neutrophils led to a decrease in neutrophil inflammatory functions including reduced ROS production, NETosis, migration and cytokine release, which they described as neutrophil “stunning”.^45^ Contrary to their published data, we identify that neutrophil functions are enhanced, and not suppressed. In addition, the depletion of tissue-resident AMØs occurs prior to O_3_ exposure and appears to have a direct impact on efferocytosis, independent of neutrophil function. This suggests that “stunning” of neutrophils is not driving our O_3_ phenotype. Additionally, recent data from Akalu et al.^46^ identified that commercially available MerTK^-/-^ mouse lines are not specific to MerTK and also delete non-MerTK gene sequences. Using a newly developed MerTK-null mouse, they demonstrated that some phenotypes previously ascribed to MerTK are no longer observed, specifically non-efferocytosis phenotypes. They did identify that efferocytosis effects appeared consistent across the various MerTK^-/-^ strains. Since our focus was on efferocytosis, we feel the use of our MerTK strain to assess O_3_-exposure responses is justified.

In summary, our results show that monocyte-derived AMØs are not recruited following O_3_ exposure. We further define that tissue-resident AMØs function in clearance of apoptotic neutrophils, which facilitates the resolution of O_3_-induced lung inflammation via MerTK. This study defines the importance of tissue-resident AMØ-mediated efferocytosis in mild resolving lung injury. Furthermore, it highlights the importance of defining distinct roles of AMØ subtypes as a means of understanding their distinct roles in lung injury and its resolution.

## Acknowledgements

The authors acknowledge contributions of the Duke Cancer Institute Flow Cytometry Facility and the Rodent Inhalation Core at Duke University, Durham, NC. Flow cytometric analysis of the human BAL specimens was performed at Northwestern University Flow Cytometry Core Facility. Northwestern University Flow Cytometry Core Facility is supported by NCI Cancer Center Support Grant P30 CA060553 awarded to the Robert H. Lurie Comprehensive Cancer Center. Cell sorting was performed on a BD FACSAria SORP cell sorter purchased through the support of NIH 1S10OD011996-01.

