## Supplemental Methods, Figures, and Tables for "Tissue-resident alveolar macrophages reduce O_3_-induced inflammation via MerTK mediated efferocytosis"

*Mouse bronchoalveolar lavage and measures of inflammation and injury*

Bronchoalveolar lavage (BAL) was performed as outlined in prior publications.[1,2] In brief, for BAL, mice were deeply anesthetized with an intraperitoneal injection of ketamine (100 mg/kg), xylazine (100 mg/kg), and saline (0.9%), dosed by weight (~350-500 µL/mouse). Following anesthesia, the chest and trachea were exposed by dissection and a small nick was made in the trachea. PE-60 tubing (Clay Adams) was used to cannulate the trachea. The tubing was connected to 12-inch tubing from an infusion set (Terumo). PBS was infused via the tubing to 20 cm $H_2O$ via connection to syringe on a ring stand until the lung reached total lung capacity. The fluid was then passively drained and the volume was recorded. The BAL cells were isolated from the fluid by centrifugation, (Eppendorf centrifuge 5424 R) (1,500 rpm, 10 min, 4ºC). BAL fluid was decanted and used for measures of lung inflammation and injury including neutrophil elastase (cat#: ab252356) and multiplex ELISA for cytokines/chemokines (#cat: PPX-08-MX47XMC) following the manufacture's protocols. To enumerate BAL cells, cells underwent red blood cell lysis with RBC lysis buffer (Biolegend, #420302) and the number of live cells were determined using a Cellometer K2 (Nexcelom Bioscience). Following total cell count, cells were immobilized by cytospin, stained with Diff-Quik (Fisher Scientific), and counted in a blinded fashion to determine exposure changes in cell differentials.

*Mouse lung tissue harvest and flow cytometry*

Flow cytometry was performed on whole lung tissue using a previously published protocol with modifications, as described below.[1,3] In brief, mice were pre-treated subcutaneously with 50µL of 1000/mL heparin. Approximately 10min after the injection, the mice were euthanized with isoflurane. The chest wall was opened, and the trachea cannulated. Following cardiac perfusion with 10 mL of PBS to remove intravascular red cells, the lung was insufflated just past total lung capacity with a digestion solution (1.5mg/mL of Collagenase A [Roche] and 0.4mg/mL DNase I [Roche] in HBSS plus 5% fetal bovine serum and 10mM HEPES). The lung tissue was excised and then incubated at 37°C for 30 min in the digestion solution with intermittent vortexing. Enzyme activity was halted by mixing the digestion mix with cold PBS then the mixture passed through a 70 µM cell strainer (Olympus Plastics, Genesee Scientific) and centrifuged (350 x g for 6-8 min at 4$^{o}$C). Cell pellets were resuspended in RBC lysis buffer and incubated on ice and washed per manufacturer's instructions for 2-3 cycles as needed (RBC Lysis Buffer, Biolegend). The cells were resuspended in cold PBS and kept on ice while aliquots were counted then apportioned for live/dead staining (Zombie UV, Biolegend) followed by cell surface staining with an optimized antibody panel (Supplemental Table 1). Stained cells were fixed (0.4% paraformaldehyde in PBS) for later acquisition (within 1 week of initial harvest). Lung cells from control animals (C57Bl/6J) were stained similarly & used for single-color controls. See Supplemental Table 1 for antibody list. Data was collected on a BD LSR Fortessa X-20 using the configuration noted in Supplemental Table 2 and analyzed using FlowJo™ version 10 software (BD Life Sciences).

*Human laboratory exposure to filtered air or ozone*

Human exposure studies were performed after informed consent through Duke Institutional Review Board approved protocols (Pro00088966 and Pro00100375). Healthy human subjects were recruited for inclusion in this study through advertisements. No study procedures were performed before obtaining informed consent. Requirements for inclusion included normal range of body mass index, no evidence of respiratory disease, and non- smoking. Exclusion criteria included a recent respiratory infection (within 4 weeks), history of smoking, pregnancy, ages less than 18 or greater than 50 years, and BMI> 35.0 and failure to understand the study protocol. Additionally, participants could not be taking antihistamines, non-steroidal anti-inflammatory drugs, or supplemental vitamins for one week prior to the study, and throughout the study's duration.

Subject exposures to filtered air or ozone were performed in a custom designed 730 cubic feet exposure room that included an HEPA air ventilation system to remove particulates. The air supply was set to a selected range of temperatures (20-23°C) and relative humidity (45-22 55%). Exposures lasted 135 minutes in duration, during which participants alternated between resting and walking on a treadmill at 2-3 mph to mimic an individual performing mildly strenuous activity under ambient conditions. Ozone exposure consisted of a dose of 200 ppb and was continuously monitored. This concentration of ozone exposure exceeds the federal government's standard (80 ppb for 8-hour average) but is similar to ambient ozone levels in the Research Triangle area during ozone alert days. Ozone was created from a 100% $O_2$ source by cold plasma corona discharge (Ozotech, Yreka CA), and mixed with filtered air before addition to chamber. The order of filtered air or ozone exposure was randomized for every participant, with at least a 21-day washout period between exposures.

*Human bronchoscopy and bronchoalveolar lavage (BAL)*

On the day following filtered air or ozone exposure, approximately 21 hours after exposure, participants underwent a flexible bronchoscopy with bronchoalveolar lavage. Subjects received sedation with midazolam (Versed) and fentanyl (Sublimaze), and topical anesthetic lidocaine prior to bronchoscopy. A flexible bronchoscope was inserted through the mouth, past the vocal cords, into the lungs. The bronchoscope was wedged into the right middle lobe and 150 mL of sterile saline was discharged into the lungs and subsequently removed with gentle syringe suction in order to obtain bronchiole alveolar lavage fluid (BALF). BALF was passed a strainer and 10mL of the sample are separated. The sample was then mixed 1:1 with HypoThermasol FRS (StemCell Technologies, USA) and then sent overnight to Northwestern University for flow cytometry and cell sorting.

*Human flow cytometry*

BAL fluid samples were filtered through a 70μm cell strainer, pelleted by centrifugation at 400rcf for 10 min at 4°C, followed by hypotonic lysis of red blood cells with 2 mL of PharmLyse (BD Biosciences) reagent for 2 minutes. Lysis was stopped by adding 13 mL of MACS buffer (Miltenyi Biotech). Cells were pelleted again and resuspended in 100 μL of 1:10 dilution of Human TruStain FcX (Biolegend) in MACS buffer, and a 10 μL aliquot was taken for counting using K2 Cellometer (Nexcelom) with AO/PI reagent. The cell suspension volume was adjusted so the concentration of cells was always less than $5x10^7$ cells/mL and the fluorophore-conjugated antibody cocktail was added in 1:1 ratio (Supplemental Table 3). After incubation at 4°C for 30 minutes, cells were washed with 5 mL of MACS buffer, pelleted by centrifugation, and re-suspended in 500 μL of MACS buffer with 2 μL of SYTOX Green viability dye

(ThermoFisher). Cells were analyzed by a FACS Aria III SORP instrument. Sample processing was performed in BSL-2 facility using BSL-3 practices. Analysis of the flow cytometry data was performed using FlowJo 10.6.2. using uniform sequential gating strategy reported in a previous publication by the Misharin group[5] and reviewed by two investigators (SS, AVM). Relative cell type abundance was calculated as a percentage out of all singlets/live/CD45+ cells.

*Efferocytosis assay*

Immortalized human T lymphocytes (Jurkat cells) (ATCC) were stained with Calcein AM (20 uL dye per 30 million cells) and incubated for 1.5h at 37$^{o}$C. The cells were then recounted and reconstituted in Jurkat cell complete media (RPMI1640 + GlutaMax, Gibco #61870-036, 10% FBS HI, 5mL A/A) at 1.5 million cells per 10 cm dish. The cells were then irradiated in the UV Stratalinker 1800 at 400 μjoules (10-15sec). The following day the cells were spun down (1,200rpm X 6min) and re-suspended at a concentration of 4 million cells in 50 uL in PBS. Jurkat cells (4 million cells, in 50 uL of PBS) or PBS control were instilled in C57Bl/6J mice by oropharyngeal aspiration for 1.5h prior to the harvest.[6] Following the harvest, BAL fluid was collected and single color flow cytometry was performed by flow cytometry to define the number of Calcein AM-positive cells. Data was expressed as a ratio of the number of recovered Jurkat cells versus the total number of Calcein AM positive Jurkat cells initially instilled.

*Real-Time PCR*

Total RNA was collected utilizing RNeasy Plus Mini Kit (Qiagen) per manufacturer protocol. RNA samples were then reverse transcribed into cDNA using Maxima H Minus cDNA Synthesis Master Mix (thermos scientific). PCR amplification was completed using the following program: 20 uL reaction volume; 50$^{o}$C, 2min; 95$^{o}$C, 2min; 40 cycles, 95$^{o}$C 1 sec, 60$^{o}$C 30 sec. All real time quantitative PCR reactions were completed using the Quant Studio 6 Flex (Applied Biosystems). The reactions utilized SyBR Green reagent (Applied Biosystems), sterile UltraPure Distilled water (Invitrogen), and primers, as listed below. MerTK forward (Sigma-Aldrich), 5'-CTGGATATTAGATGGACGAAG-3', MerTK reverse (Sigma-Aldrich), 5'-AGAAAGCTCTTTGTAAGTCC-3', 18S forward (Integrated DNA Technologies), 5'-TTGACGGAAGGGCACCACCAG-3', 18S reverse (Integrated DNA Technologies), and 5'-GCACCACCACCCACGGAATCG-3'. Gene expression values were normalized to 18S as a housekeeper and presented as a fold change normalized again to FA.

| Mouse Target | Clone | Isotype | Conjugate | Working Dilution | Vendor | Catalogue # |
|---|---|---|---|---|---|---|
| **CD31 (PECAM-1)** | MEC13.3 | Rat IgG2a, κ | PerCP-Cy5.5 | 1:400 | Biolegend | 102522 |
| **Ly6C** | HK1.4 | Rat IgG2a, κ | PerCP-Cy5.5 | 1:200 | Biolegend | 128012 |
| **B220 (CD45R)** | RA3-6B2 | Rat IgG2a, κ | PE | 1:100 | Biolegend | 103208 |
| **CD49b** | DX5 | Rat IgM, κ | PE | 1:100 | Biolegend | 108908 |
| **Siglec-F (CD170)** | E50-2440 | Rat IgG2a, κ | PE-CF594 | 1:1500 | BD Horizon™ | 562757 |
| **F4/80 (Ly71)** | BM8 | Rat IgG2a, κ | PE-Cy7 | 1:400 | Biolegend | 123114 |
| **CD163** | S15049I | Rat IgG2a, κ | APC | 1:200 | Biolegend | 155306 |
| **CD169 (Siglec-1)** | 3D6.112 | Rat IgG2a, κ | APC | 1:100 | Biolegend | 142418 |
| **CD206** | C068C2 | Rat IgG2a, κ | APC | 1:100 | Biolegend | 141708 |
| **MERTK (Mer)** | 2B10C42 | Rat IgG2a, κ | APC | 1:50 | Biolegend | 151508 |
| **Isotype Control** | RTK2758 | Rat IgG2a, κ | APC | 1:50 | Biolegend | 400512 |
| **Ly6G** | 1A8 | Rat IgG2a, κ | AF700 | 1:200 | Biolegend | 127622 |
| **CD11b** | M1/70 | Rat IgG2a, κ | APC-Cy7 | 1:150 | Biolegend | 101226 |
| **CD64 (FcγRI)** | X54-5/7.1 | Rat IgG2a, κ | BV421 | 1:50 | Biolegend | 139309 |

| | | | | | | |
|---|---|---|---|---|---|---|
| **CD3ε** | 145-2C11 | Rat IgG2a, κ | BV510 | 1:100 | Biolegend | 100353 |
| **CD103** | 2E7 | Rat IgG2a, κ | BV510 | 1:100 | Biolegend | 121423 |
| **CD45** | 30-F11 | Rat IgG2a, κ | BV605 | 1:500 | Biolegend | 103155 |
| **I-A/I-E** | M5/114.15.2 | Rat IgG2b, κ | BV650 | 1:1500 | Biolegend | 107641 |
| **CD24** | M1/69 | Rat IgG2b, κ | BV711 | 1:800 | Biolegend | 563450 |
| **CD11c** | N418 | AH IgG | BV785 | 1:100 | Biolegend | 117336 |
| **CD8_BUV395** | 53-6.7 | Rat IgG2a, κ | | 1:100 | BD Horizon™ | 563786 |
| **Zombie UV™** | | Fixable viability dye | 350ex / 459em | 1:1000 | Biolegend | 423108 |
| **CD4_BUV805** | GK1.5 | Rat IgG2b, κ | | 1:100 | BD Horizon™ | 612900 |

Supplemental Table 1. Immunophenotyping: Mouse: Antibodies and Staining reagents.

| Detector | PMT | Mirror/ Beam Splitter | Band Pass Filter | Fluorochrome |
|---|---|---|---|---|
| **770LP_820_60** | A | 770 | 820_60 | BUV805 |
| **450LP_515_30** | B | 450 | 515_30 | Zombie UV |
| **379_28** | C | ---- | 379_28 | BUV395 |
| **750LP_780_60** | A | 750 | 780_60 | BV786 |
| **690LP_710_50** | C | 690 | 710_50 | BV711 |
| **635LP_670_30** | D | 635 | 670_30 | BV650 |
| **600LP_610_20** | E | 600 | 610_20 | BV605 |
| **495LP_525_50** | G | 495 | 525_50 | BV510 |
| **450_50** | H | ---- | 450_50 | BV421 |
| **635LP_710_50** | A | 685 | 695/40 | PerCP-Cy5.5 |
| **495LP_525_50** | B | 505 | 525_50 | GFP |
| **488_10** | C | ---- | 488_10 | Side Scatter (SSC) |
| **Diode** | FSC | | | |
| **Diode** | SSC | | | |
| **735LP_780_60** | A | 735 | 780_60 | PE-Cy7 |
| **655LP_710_50** | B | 655 | 710_50 | -- |
| **635LP_695_40** | C | 635 | 695_40 | -- |
| **595LP_610_20** | D | 595 | 610_20 | PE-CF594 |
| **575_26** | E | ---- | 575_26 | PE |
| **750LP_780_60** | A | 750 | 780_60 | APC-Cy7 |

| | | | | |
|---|---|---|---|---|
| **710LP_730_45** | B | 710 | 730_45 | AF700 |
| **660_20** | C | ---- | 670_30 | APC |

Supplemental Table 2. LSRFortessa X-20 configuration.

| **Antigen** | **Clone** | **Fluorochrome** | **Manufacturer** | **Cat #** |
|---|---|---|---|---|
| CD4 | RPA-T4 | BUV395 | BD | 564724 |
| CD19 | HIB19 | BUV395 | BD | 740287 |
| CD25 | 2A3 | BUV737 | BD | 564385 |
| CD56 | NCAM16.2 | BUV737 | BD | 612766 |
| HLA-DR | L243 | eFluor450 | ThermoFisher | 48-9952-42 |
| CD45 | HI30 | BV510 | Biolegend | 304036 |
| CD15 | HI98 | BV786 | BD | 563838 |
| Live/Dead | Not applicable | SYTOX Green | ThermoFisher | S34860 |
| CD3 | SK7 | PE | ThermoFisher | 12-0036-42 |
| CD127 | HIL-7R | PECF594 | BD | 562397 |
| CD206 | 19.2 | PECy7 | ThermoFisher | 25-2069-42 |
| CD8 | SK1 | APC | Biolegend | 344721 |
| CD14 | M5E2 | APC | Biolegend | 301808 |
| EpCAM | 9C4 | APC | Biolegend | 324208 |

Supplemental Table 3.  Human: Flow cytometry panel used for BAL sample phenotyping

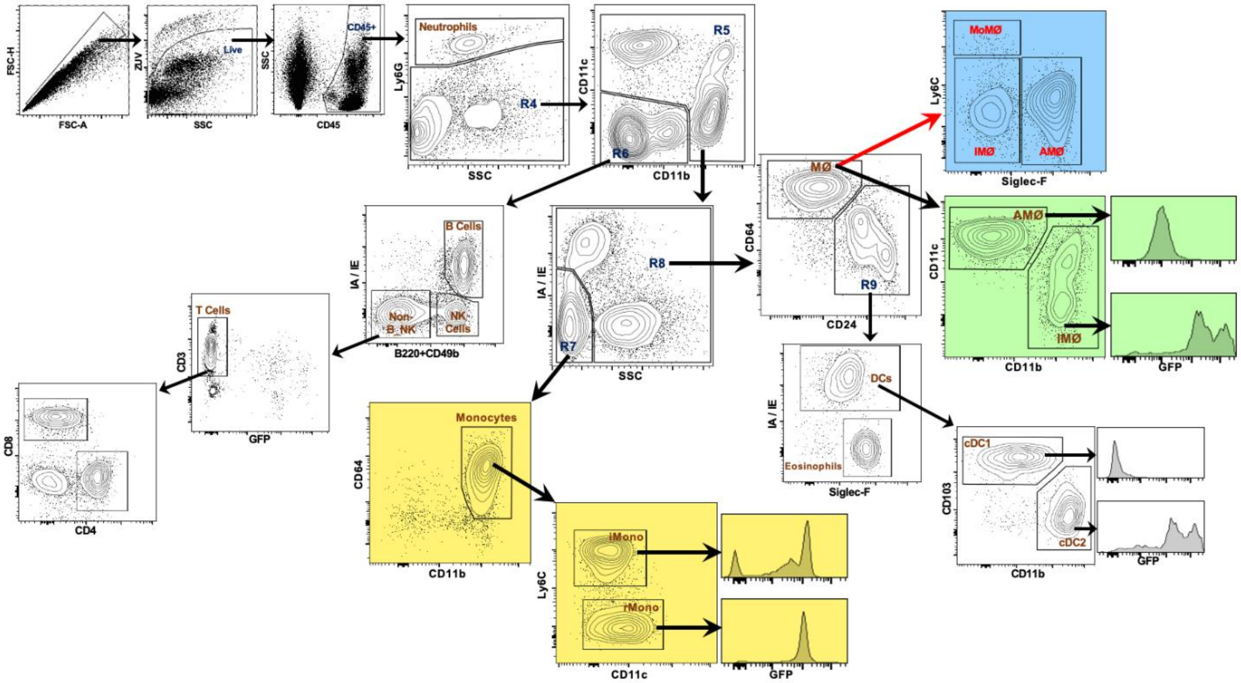


Supplemental Figure 1. **Mouse Flow Cytometry Gating Strategy.** Whole lung tissue was digested and stained for flow cytometry. Individual immune cells were defined based on specific cell surface markers based on prior published protocols (representative sample). Color gating indicates cellular and lineage tracking of specific cell populations of interest: yellow (monocytes), green (macrophages), blue (alternative macrophage gating to segregate monocyte-derived macrophages (MoMØ) from interstitial macrophages (IMØ) and alveolar macrophages (AMØ)).

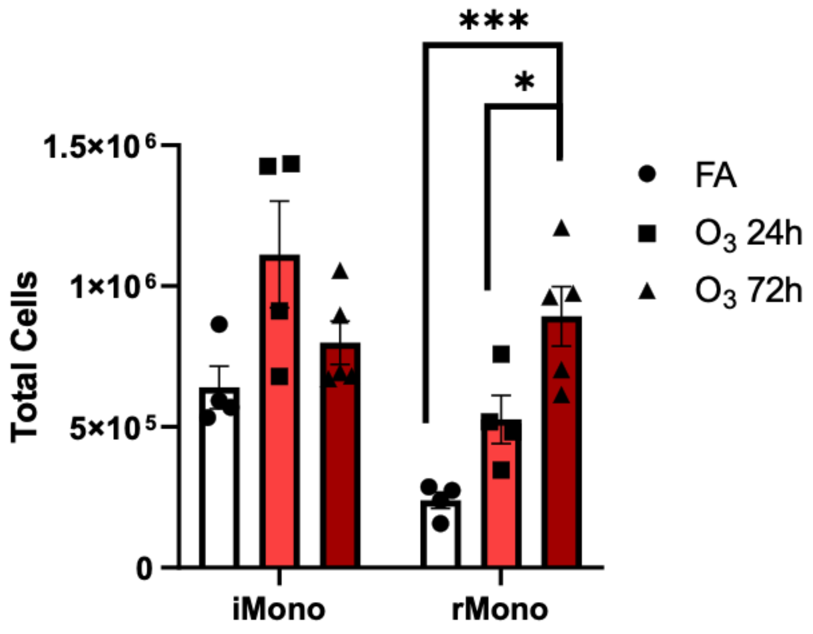


Supplemental Figure 2. **Inflammatory and constitutive monocytes are increased in lung tissue in response to acute $O_3$ exposure.** Monocyte lineage labeling was induced in Cx3cr1$^{ERCre}$ x zsGreen mice with tamoxifen. Lineage label was induced with tamoxifen 24h prior to exposure with filtered air or $O_3$ (2 ppm) for 3h. 24 or 72h post exposure whole lungs were processed, stained and analyzed by flow cytometry. Following $O_3$ exposure GFP$^+$ inflammatory (iMono) and constitutive monocytes (rMono) were increased. n=4-5 mice per group and replicated x 1. *$p<0.05$. ANOVA analysis was conducted using Tukey's Honestly Significant Difference Test post-hoc.

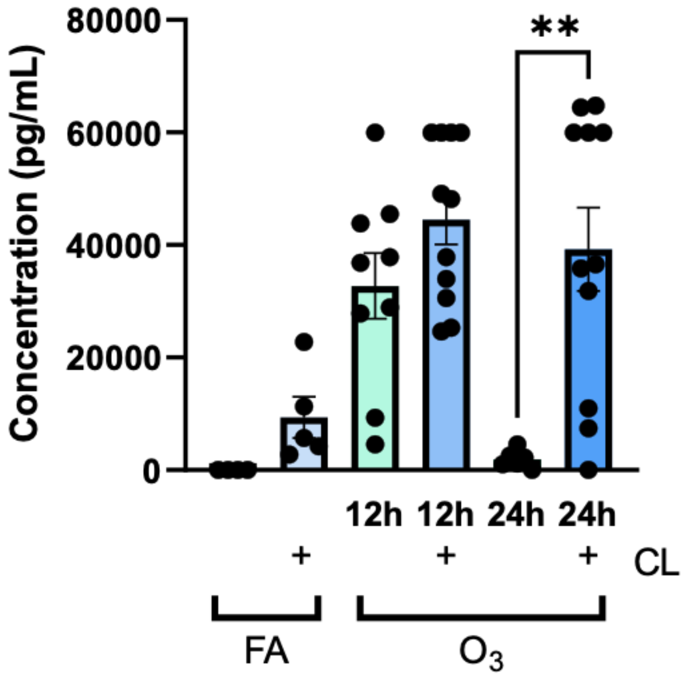


Supplemental Figure 3. **Neutrophil elastase is persistently elevated in tissue-resident AMØ depleted mice.** PBS or Clodronate (CL) was administered to male C57BL/6J mice 72h prior to FA or $O_3$ (2 ppm) for 3h. Mice were harvested at 12 and 24h post exposure and BAL was collected and processed. Bars shaded green indicate PBS/vehicle control while blue shades indicate CL administration. BAL fluid from PBS or CL and FA/O3 exposed mice was assessed for neutrophil elastase concentration. n=4-10 mice per group/exposure/timepoint *p<0.05. ANOVA analysis was conducted using Tukey's Honestly Significant Difference Test post-hoc.

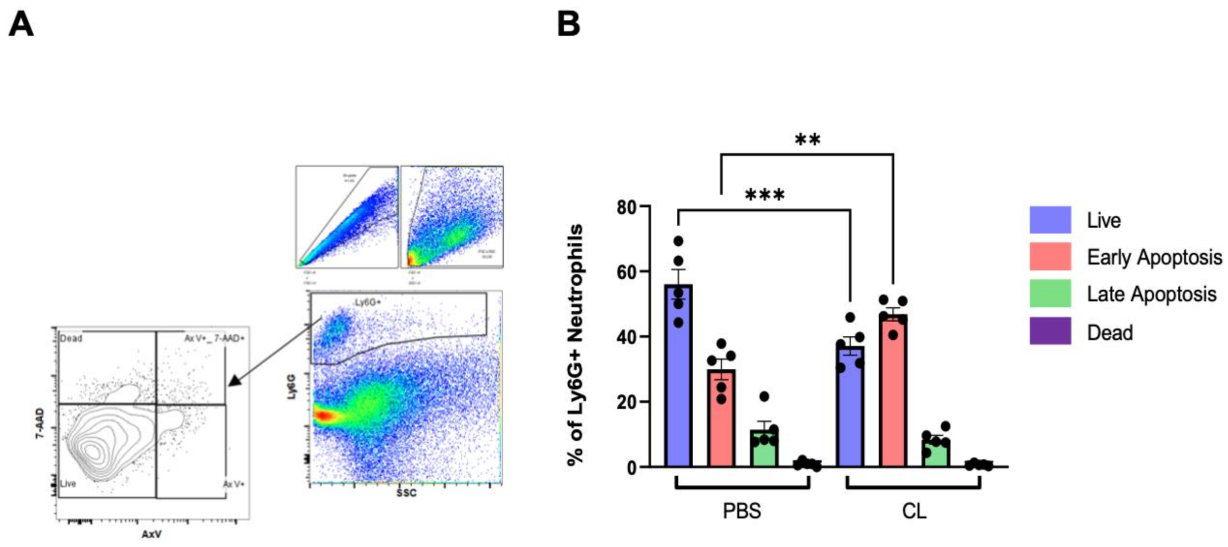


Supplemental Figure 4. **Tissue-resident AMØ depletion increased the proportion of apoptotic neutrophils following $O_3$ exposure.** A. The gating strategy to delineate neutrophil populations. Flow plots are a representative example of n=5 samples per condition. B. PBS or Clodronate (CL) was administered to male C57BL/6J mice 72h prior to $O_3$ (2 ppm) for 3h. 24h post-exposure, BAL was collected, stained with Ly6G, 7AAD and Annexin V and analyzed by flow cytometry. Neutrophils were defined as Ly6G$^+$ cells and then assessed for 7AAD and Annexin V staining. “Live” neutrophils were defined as Ly6G$^+$, 7AAD$^-$, Annexin V$^-$, “Early Apoptosis” neutrophils were defined as Ly6G$^+$, 7AAD$^-$, Annexin V$^+$, “Late Apoptosis” neutrophils were defined as Ly6G$^+$, 7AAD$^+$, Annexin V$^+$, and “Dead” neutrophils were categorized as Ly6G$^+$, 7AAD$^+$, Annexin V$^-$. **$p<0.005$, ***$p<0.0005$
